# Evolutionary changes of phospho*enol*pyruvate transporter (PPT) during the emergence of C_4_ photosynthesis

**DOI:** 10.1101/713537

**Authors:** Ming-Ju Amy Lyu, Yaling Wang, Jianjun Jiang, Genyun Chen, Xin-Guang Zhu

## Abstract

C_4_ photosynthesis is a complex trait, which evolved from its ancestral C_3_ photosynthesis by recruiting pre-existing genes. The evolutionary history of enzymes involved in the C_4_ shuttle has been extensively studied. Here we analyze the evolutionary changes of phospho*enol*pyruvate (PEP) transporter (PPT) during its recruitment from C_3_ to C_4_ photosynthesis. Our analysis shows that 1) among the two PPT paralogs, i.e. PPT1 and PPT2, PPT1 is an ancestral copy while PPT2 is a derived copy; 2) during C_4_ evolution, PPT1 shifted its expression from predominantly in root to in leaf, and from bundle sheath cell to mesophyll cell, supporting that PPT1 was recruited into C_4_ photosynthesis; 3) PPT1 gained increased transcript abundance, gained more rapid and long-lasting responses to light during C_3_ to C_4_ evolution, while PPT2 lost its responsiveness to light; 4) PPT1 gained a number of new *cis*-elements during C_4_ evolution; 5) C_4_ PPT1 can complement the phenotype of *Arabidopsis* PPT1 loss-of-function mutant, suggesting that it is a bidirectional transporter and its transport direction did not alter during C_4_ evolution. We finally discuss mechanistic linkages between these observed changes in PPT1 and C_4_ photosynthesis evolution.

**High light:** During the process of C_4_ photosynthesis evolution, PPT not only experienced changes in its expression location and transcript abundance, but also acquired new *cis*-elements in its promoter region and accumulated protein variations.

## Introduction

Compared to C_3_ photosynthesis, C_4_ photosynthesis has higher light, nitrogen and water using efficiencies (Sage and Zhu, 2011). It achieves these superior traits through a CO_2_ concentrating mechanism operating in a special leaf anatomical feature termed as “Kranz anatomy” (Hatch, 1987). The CO_2_ concentrating mechanism involves many enzymes and metabolite transporters, which together pumps CO_2_ from mesophyll cell (MC) to bundle sheath cell (BSC), creating a local high CO_2_ environment in the BSC where the Calvin Benson cycle operates (Hatch and Osmond, 1976). Though C_4_ photosynthesis shows greater light using efficiency than C_3_ photosynthesis, all genes required for the operation of C_4_ photosynthesis pre-existed in its C_3_ ancestors and play crucial house-keeping functions (Aubry *et al.*, 2011). The evolution of C_4_ photosynthesis therefore represents an outstanding example of recruitment and re-organization of plant primary metabolism to fulfill new functions (Burgess *et al.*, 2016b; West-Eberhard *et al.*, 2011).

Most of the known enzymes and transporters involved in C_4_ photosynthesis are encoded by multigene families (Moreno-Villena *et al.*, 2018) and usually only one of these paralogs is recruited to perform C_4_ function. So far, the evolutionary recruitment of most of the C_4_ shuttle enzymes, such as phospho*enol*pyvuvate carboxlase (PEPC) (Christin and Besnard, 2009; Christin *et al.*, 2013; Moreno-Villena *et al.*, 2018), phospho*enol*pyvuvate carboxykinase (PEP-CK) (Christin *et al.*, 2013; Christin *et al.*, 2009; Moreno-Villena *et al.*, 2018), NADP-malic enzyme (NADP-ME) (Christin *et al.*, 2013; Moreno-Villena *et al.*, 2018), pyruvate phosphate kinase (PPDK) (Christin *et al.*, 2013; Moreno-Villena *et al.*, 2018), and malate dehydrogenase (MDH) (Christin *et al.*, 2013; Moreno-Villena *et al.*, 2018) have been intensively studied. These studies show a number of properties associated with the recruited paralogs for C_4_ function, which include relatively high expression levels (Moreno-Villena *et al.*, 2018), suitable enzyme catalytic property mainly through accumulated mutations in the coding region (Christin *et al.*, 2013), and being regulated by light and chloroplast-to-nucleus signals (Reyna-Llorens and Hibberd, 2017).

So far, compared to the relatively intense study of the evolution of C_4_ shuttle enzymes, the evolution of metabolite transporters is less studied. Compared to C_3_ photosynthesis, the extensive usage of transporters is a major feature of C_4_ photosynthesis (Weber and von Caemmerer, 2010). In fact, to produce one molecule of triose phosphate for the synthesis of sucrose, only one transporter is needed in C_3_ photosynthesis, while at least 30 metabolite transport steps are involved in NADP-ME type C_4_ photosynthesis (Weber and von Caemmerer, 2010). Furthermore, the flux through the transporters is much higher in C_4_ photosynthesis. This is because different from C_3_ photosynthesis, where the end-product of photosynthesis, i.e. triose phosphate (TP), is exported as one unit, i.e. the flux through triose phosphate transporter is 1/3 of the photosynthetic CO_2_ uptake rate, during the operation of C_4_ photosynthesis, the flux of the metabolite transport between different compartments is higher than the photosynthetic CO_2_ uptake rate due to the leakage of CO_2_ from BSC to MC. This, together with the typically higher photosynthetic CO_2_ uptake rate in C_4_ leaves compared to C_3_ leaves, C_4_ photosynthesis demands a much higher capacity for metabolite transport between the two compartments (Hatch and Osmond, 1976; von Caemmerer and Furbank, 2003). Indeed, a number of transporters on chloroplast envelope, including PEP transporter (PPT), pyruvate transporter (BASS2), and malate transporter in MC (DIT1), all show much higher transcript abundance in C_4_ species than in C_3_ species (Emms *et al.*, 2016; Lyu *et al.*, 2018; Moreno-Villena *et al.*, 2018). Identifying the C_4_ paralogs of individual metabolite transporters, understanding their evolutionary trajectories and the molecular mechanisms behind the increased abundance or capacity of these transporters are major research focuses of the current C_4_ photosynthesis research.

In this study, we aim to characterize the evolutionary changes of the transporter of PEP (PPT), a critical metabolite transporter involved in C_4_ photosynthesis as PEP is the substrate for PEPC and its carboxylation underlies the first step of the C_4_ acid formation in C_4_ photosynthesis. We first constructed phylogenetic tree of PPT orthologs and showed that PPT1 is the ancestral copy while the PPT2 is the derived copy. We recapitulated that PPT1, the version recruited to operate in C_4_ photosynthesis, showed increased transcript abundance, shifted its cell specificity of expression from root to leaf, and from BSC to MC. We further showed that PPT1 gained more rapid and long-lasting responses to light in C_4_ as compared to the closely related C_3_ species. We further predicted a set of candidate *cis-*elements that are potentially responsible for these newly acquired expression features of PPT1 in C_4_ photosynthesis. Finally, we identified amino acid modifications during the C_3_ to C_4_ transition in *Flaveria* and thus suggested that the PPT1 in C_4_ species is a bi-directional transporter using transgenetic experiments. All these findings are discussed in light of the role of PPT in coping with stresses plant faced during the evolution from C_3_ to C_4_ photosynthesis.

## Materials and Methods

### Construction of the PPT phylogenetic tree

To construct the phylogenetic tree of PPT, we used protein sequences from 23 species, which included representative species along the phylogeny of viridiplantae, spanning from basal species belonging to Chlorophyte (*Micromonas.pusilla* and *Chlamydomonas reihardtii*), to Embryophyte (*Marchantia.polymorpha*), to Tracheophyte (*Selaginella.moellendorfii*), and to species belonging to Angiosperm (*Amborella trichopoda*), including 10 species belonging to eudicot and 8 species belonging to monocot (Fig. 1). For the eudicot, we included 6 species of Brasssicadeae; in the monocot, we included 7 species belong to the grass family (Fig. 1).

**Figure 1.**
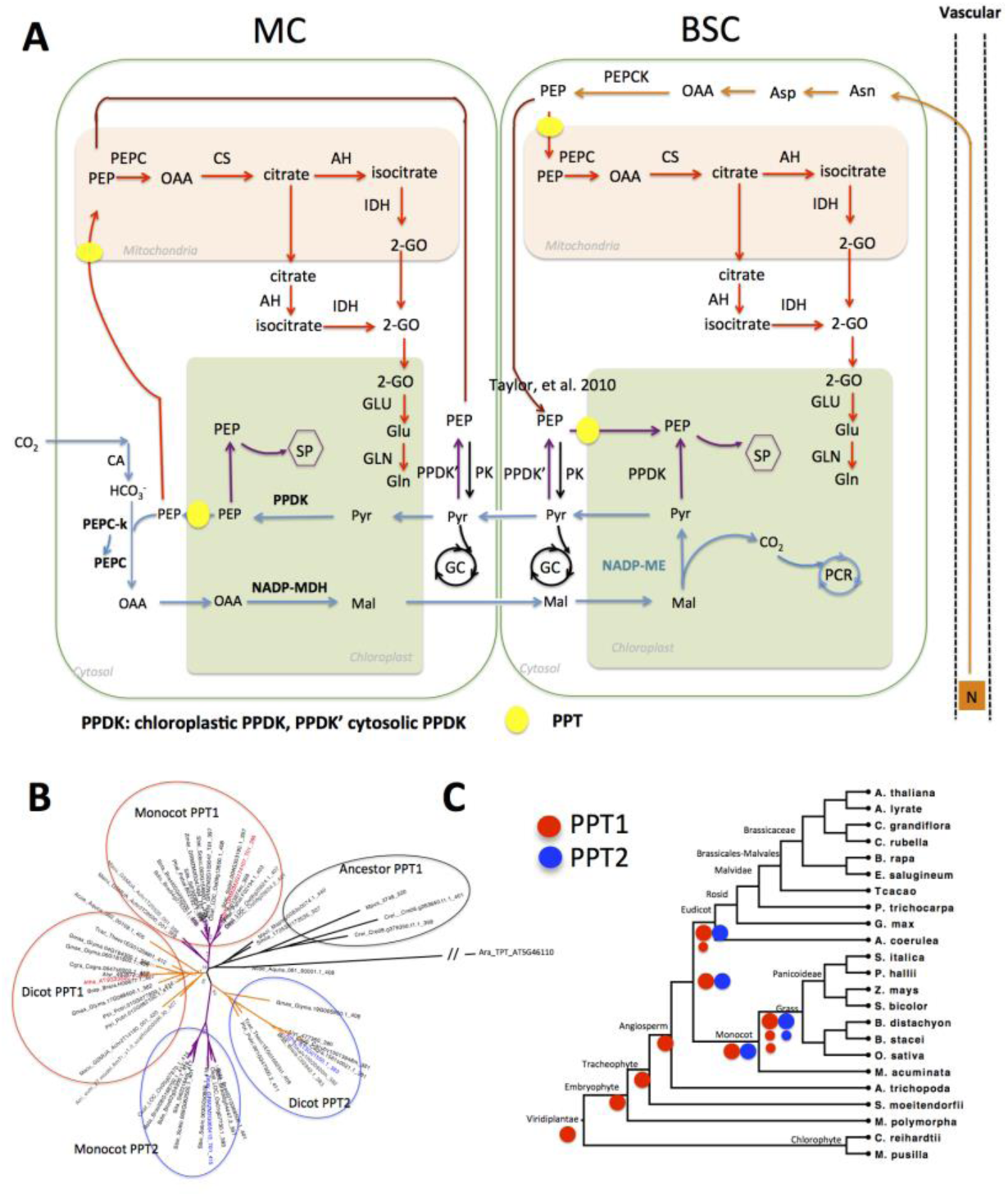
The function and evolution of PPT. (A) shows PEP-related metabolic pathways in C_4_ *Flaveria* species. In MC, PEP is used as substrate for PEPC catalyzed carboxylation (blue lines). It is also a substrate of Shikimate Pathway (SP) in chloroplast, which is expected to exist in both MC and BSC (purple lines); moreover, PEP is a substrate for citrate pathway in mitochondrion (red lines) and glycolysis in cytosol (black lines). PEP is also involved in nitrogen recycle from xylem (orange lines). (B) The gene tree of PPT family genes from 23 representative species of Viridiplantae. The tree was inferred from the alignment of protein sequence of PPT based on maximum likelihood method. The numbers besides each node are the bootstrap scores from 1000 bootstrap sampling. (C) Schematic depiction of the evolution of PPT1 and PPT2 based on phylogenetic relationship of species. PPT1 has one or two copies in eudicot species and two or three copies in monocot species. Red circles represent PPT1 and blue circles represent PPT2, large circles strand for original copy and smaller circles stand for duplicated copies after the division of monocot and dicot. The phylogenetic relationship of species was inferred from the Phytozome website. (Abbreviations: MC: mesophyll cell; BSC, bundle sheath cell; SP, shikimate pathway, PCR: photosynthetic carbon reduction, GC: glycolysis cycle.)

The genome-wide protein sequences of these 23 species were downloaded from Phytozome (http://phytozome.jgi.doe.gov/). We used the protein sequences of PPT1 (AT5G33320) and PPT2 (AT3G01550) from *Arabidopsis thaliana* (*Arabidopsis*) as query sequences to search orthologs in other 22 species using the blastp algorithm in BLAST+ (v2.2.31) package (Camacho *et al.*, 2009), with a threshold of blast score >= 250 and evalue <1E-5. All orthologs sequences were then aligned using MUSCLE (Edgar, 2004) with default parameters. Gene trees were constructed using the RAXML software (Stamatakis, 2006) based on protein sequence alignment using the PROTGAMMAILG model. The robustness of the tree topology was evaluated by bootstrap scores, which was calculated from 1000 independently constructed gene trees.

### Surveying the transcript abundance of PPTs from published RNA-SEQ data

Since the transcript abundance and expression location can provide insights for the function of a gene, we further compared the transcript abundance of PPT1 and PPT2 among different photosynthetic types of species, and their expression patterns in tissues and cell types using publicly available RNA-SEQ data. We surveyed RNA-SEQ data from four independent C_4_ lineages, namely: *Heliotropium*, *Mollugo*, *Neurachne* and *Flaveria* available from 1KP (http://www.onekp.com/blast.html). The RNA-SEQ source and quantification process were documented in our previous work (Lyu *et al.*, 2018). When comparing transcript abundance of PPTs in roots and leaves from C_3_ and C_4_ species, we surveyed RNA-SEQ data from two *Flaveria* species (Lyu *et al.*, 2018), two *Brassicacae* species (*G. gyanndara* and *T. hassleriana*) (Kulahoglu *et al.*, 2014), and 21 species in the grass family (Moreno-Villena *et al.*, 2018). We also compared the transcript abundance of PPTs in BSC and whole leaf or MC in both C_3_ species and C_4_ species. Specifically, transcript data of PPTs in BSC and whole leaf of *Arabidopsis* were from (Aubry *et al.*, 2014), and transcript data in BSC and MC of maize were from (Denton *et al.*, 2017; Li *et al.*, 2010; Tausta *et al.*, 2014), and those of *G. gynandra* were from (Chang *et al.*, 2012), and those of *S. virids* were from (John *et al.*, 2014) and those of *P. virgatum* were from (Rao *et al.*, 2016). The photosynthetic type and abbreviation of species were listed in Table S1.

### Quantification of the light response of PPTs in *Flaveria* species using qRT-PCR

*F. robusta* and *F. ramosissima*, *F. palmeri* and *F. bidentis* plants are gifts from Prof. Peter Westhoff (Heinrich-Heine-University); *F. sonorensis*, *F. australasica*, *F. trinervia*, *F. kochiana*, *F. vaganita* are gifts from Prof. Rowan Sages (University of Toronto). *Flaveria* plants were grown in soil in growth rooms with air temperature controlled at 25°C, the relative humidity at 60%, and photoperiod being 16/8 hour day/night, and PPFD being 500 μmol m^−2^ s^−1^. The *Flaveria* plants were watered twice a week and fertilized weekly. To study the gene expression differences of PPT1 and PPT2 in responses to illumination, one-month old plants were put to dark at 6 pm. The dark-adapted plants were illuminated on 9:30 am the next day. A fully expanded leaf, usually the 2^nd^ leaf pair or the 3^rd^ leaf pair counted from the top, were cut and quickly put to liquid nitrogen after the leaf was illuminated for 0.5h, 2h and four 4h respectively. Leaf samples were stored in −80 °C before processing.

RNA was extracted following the protocol of the PureLInk^TM^ RNA Mini (ThermoFisher Scientific). For qRT-PCR, 0.2-0.5 μg RNA was incubated with Superscript II Reverse Transcriptase (TransGen Biotech, Beijing). qRT-PCR was conducted following the manufacturer’s instructions of the kit UNICONTM qPCR SYBR Green Master Mix (YEASEM, Shanghai). cDNA, buffer and enzyme were pipetted to the Hard-Shell PCR Plates 96-well (BIO-RAD), and covered by Microseal ‘B’ seal (Bio-Rad). qRT-PCR was run in the BIO-RAD CFX connect system. Relative transcript abundance was calculated by comparing to ACTIN7 and data was processed using BIO-RAD CFX Maestro software. For each gene, three technical and three biological replicates were performed. The primers used here are listed in Table S2.

The promoter sequence of PPT1 and PPT2 from four species, namely, *F. robusta*, *F. sonorensis*, *F. ramosissima* and *F. trinvervia* were referenced from the draft genome sequence of the four species (unpublished), and verified by sequencing. The primers used here were listed in Table S2.

### Protein sequences of PPT1 and PPT2

The protein sequences of PPT1 and PPT2 in species from the *Flaveria* genus were predicted based on transcript sequence as described in (Lyu *et al.*, 2018). Protein sequences of orthologs were aligned using the tool MUSCLE (Edgar, 2004). We further identified consistent amino acid modifications between C_3_ and C_4_ species, which were defined as sites that show difference between C_3_ and C_4_ species, but they are conserved within C_3_ species and also conserved within C_4_ species. These identified consistent modifications were mapped to the phylogenetic tree of *Flaveria* (Lyu *et al.*, 2015) to identify the evolutionary stage of their appearance during C_4_ evolution in the *Flaveria* genus. With the protein sequence information, we also predicted the 3-D protein structure of PPT1 of *Flaveria* species using the I-TASSER online server (Yang and Zhang, 2015). Positive selection test was performed using PAML package (V4.8) (Yang, 2007) following (Huang *et al.*, 2017). To investigate the copy number of 13-aa elements in *Flaveria* species, DNA was extracted from the 2^nd^ pair or 3^rd^ pair of leaf counted from the top following the protocol of TIANquick Midi Purificatioin kit (TIANGEN Biotech, Beijing). The primers were listed in Table S2.

### Subcellular localization of *Flaveria* PPT1 and PPT2

To determine the subcellular localization of PPT1 and PPT2 from *Flaveria* species, we generated fluorescence fusion proteins by tagging a green fluorescent protein (GFP) in the C-terminus of PPT and transiently expressed it in *Nicotiana benthamiana* (tobacco) leaves. Specifically, the coding sequences (CDS) of PPT1 and PPT2 were amplified from cDNAs either reverses transcribed from RNAs of different *Flaveria* species or *de novo* synthesized DNA (*F. robusta*, by Shanghai Personalbio LLC) by PCR. A CDS with the 52aa insertion deleted, i.e. ΔFbid-PPT1, was generated via overlapping PCR. All the primers were listed in Table S2. All the PCR fragments of *PPT1* and *PPT2* were integrated into the binary vector pCAMBIA1302 via homologous recombination-based in-fusion cloning (GBClonart). The promoter used was a CaMV 35S promoter. The final plasmids were verified by Sanger-sequencing (Sangon Biotech, Shanghai). The verified vectors were transformed into *Agrobacterium tumefaciens* (Agrobacterium) strain GV3101 competent cells (TransGen Biotech). The Agrobacterium cells were cultured in liquid Luria-Bertani (LB) medium containing rifamycin and kanamycin and resuspended in infiltration buffer (10 mM MES pH5.7, 10 mM MgCl_2_, 200 μM acetosyringone) to OD_600_ ~1.0. The Agrobacterium were infiltrated into tobacco leaves with a syringe. After 36 to 48 h, the fluorescence signals from leaf pavement cells were examined with a confocal fluorescence microscope (Zeiss LSM880). The autofluroscence signal from chlorophyll was used as a marker for chloroplast thylakoid.

### Expression of *Flaveria* PPT1 in *Arabidopsis cue1-5* mutant

The *Arabidopsis* PPT1 mutant *cue1-5*, which is an EMS mutant harboring R81C mutation in PPT1, was ordered from NASC (Stock Number N3156). Then, we introduced different *Flaveria PPT1-GFP* driven by *35S* promoter into *cue1-5* mutant via Agrobacterium-mediated floral dipping method. Agrobacteria transformed with the binary plasmids were cultured in Luria-Bertani media at 28°C, pelleted and re-suspended in transformation buffer (50 g sucrose, 2.2 g Murashige and Skoog powder, 200 μL silwet L-77, and 10 μL 6-BA for 1L, pH 5.8 to 6.0) to OD_600_ ~1.0. The *Arabidopsis* flowers were dipped in bacteria and kept for 5 min, and then the plants were put under dark for overnight. The floral dipping process was repeated once more one week later. After maturation, the seeds were collected and screened on ½ MS agar plates containing hygromycin at a concentration of 35 mg/L. The positive T_1_ transformants were transferred to soil. The T_2_ lines were used for morphological phenotypes. The plants were grown in a growth-chamber with a long-day condition (16 light /8 dark), PPFD of ~100 μmol⋅ m^−2^⋅s^−1^, and temperature cycle of 23 °C during the day and 21°C at night.

## Results

### The evolutionary origin of PPT in the viridiplantae

PPT is a transporter of PEP (Knappe *et al.*, 2003), while PEP is involved in a number of metabolic pathways in higher plants. Figure 1 A shows the reactions that PEP is either a substrate or product in a typical NADP-ME type C_4_ leaf (Fig. 1A). Specifically, PEP is the substrate of PEPC and its carboxylation represents the first step of the CO_2_ fixation in C_4_ photosynthesis. PEP is also involved in the shikimate pathway in chloroplast, which generates aromatic amino acid and secondary metabolites (Fischer *et al.*, 1997; Herrmann and Weaver, 1999). Given that BSC contains chloroplasts in C_4_ plants (Stata *et al.*, 2016), shikimate pathway is expected to present in both MC and BSC. Moreover, PEP is also a substrate of citric acid cycle in mitochondria (Krebs, 1982). Recent report shows that PEP is involved in nitrogen recycling from xylem (Bailey and Leegood, 2016) and nitrogen mobilization from aging leaves (Taylor *et al.*, 2010).

To investigate the evolution of PPT in the viridiplantae, we constructed a phylogenetic tree for PPT orthologs from 23 species with genome sequences available from the Phytozome database (https://phytozome.jgi.doe.gov/pz/portal.html), which were selected to represent the major branching points of the viridiplantae phylogeny. The gene tree shows that there are two paralogs of PPT in most species with one being ortholog of *Arabidopsis* PPT1 and another being the ortholog of *Arabidopsis* PPT2 (Fig. 1B). The gene tree shows that PPT1 is the ancestral copy and PPT2 is the derived one. PPT1 is the original copy ever since Chlorophyte, whereas PPT2 is the latecomer, which was originated after the split of *Amborella trichopoda* from other angiosperm species (Fig. 1C). Interestingly, there is one or two copies of PPT1 and single copy of PPT2 in dicot species, whereas around two or three copies of PPT1 and one or two copies of PPT2 in grass species; which is consistent with an extra whole genome duplication (WGD) event in monocot species (Jiao *et al.*, 2014). Therefore, the ancestral PPT1, after gene duplication, gained extra copies, which might have facilitated neofunctionalization to support new functions in C_4_ photosynthesis.

### The evolution of PPTs in transcription along the emergence of C_4_ species

We further compared the functional difference between PPT1 and PPT2. First, we examined the transcript abundance of PPT1 and PPT2 in a few sets of species which are evolutionarily closely related but have different photosynthetic types. These species are from four genera with each representing an independent C_4_ lineage. Among these four genera, three of them were from dicot, i.e. *Flaveria, Heliotropium, Mollugo*, and one from monocot, i.e. *Neurachne*. The transcript abundance data for these species are shown in Fig. 2A. The RNA-SEQ data for the *Flaveria* species are from the 1,000 Plants project (1KP) (Matasci *et al.*, 2014) and (Mallmann *et al.*, 2014), both of which are from leaf samples, and have been demonstrated to be comparable (Lyu *et al.*, 2018). RNA-SEQ data for other three genera are from 1KP (Matasci *et al.*, 2014). In the analysis, data for mature leaves were used. The comparison shows that in C_3_ species, though PPT1 is an ancestral paralog, the PPT2 displays higher transcript abundance than the original copy except in the two C_3_ species in the *Heliotropiumi* genus, namely, *H. calcicola* (Hcal) and H. *karwinsky (*Hkar). The higher expression of PPT2 over PPT1 is also shown in C_3_-C_4_ species of *Mollugo* and *Neurachne*, as well as in C_3_-C_4_ species in *Heliotropium: H. filiforme* (Hfil) and *Flaveria*: *F. sononrensis* (Fson). During the transition from C_3_ to C_4_ photosynthesis, however, we observed a dramatic increase of transcript abundance in PPT1 while the transcript abundance of PPT2 remained relatively less changed (Fig. 2 A). As a result, in C_4_ species, PPT1 showed much higher expression abundance compared to PPT2 (Fig. 2 A).

**Figure 2.**
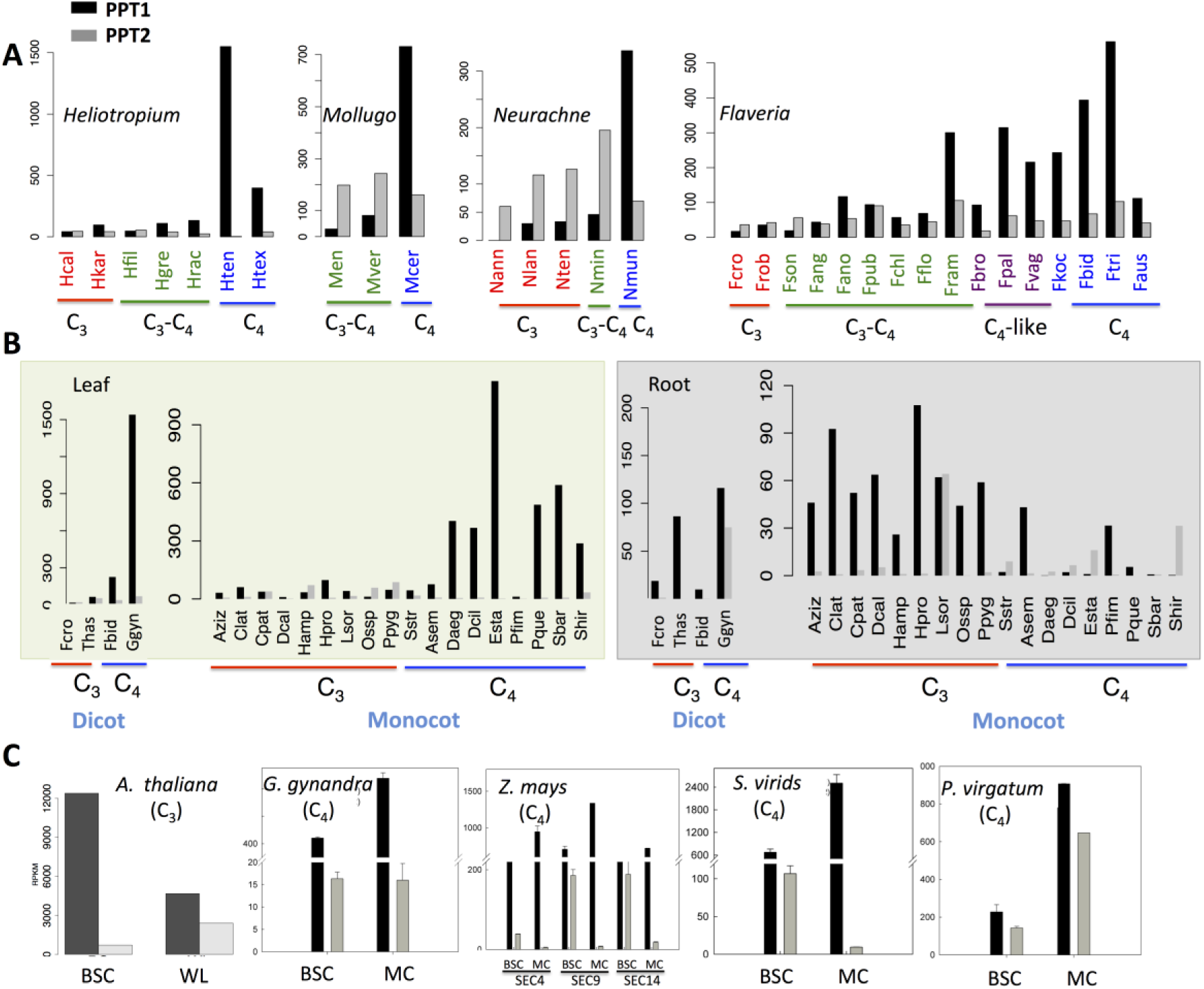
The evolution of transcript abundance of PPT1 and PPT2 from C_3_ to C_4_ species. (A) The transcript abundance of PPT1 and PPT2 along the evolution of C_4_ photosynthesis in four genera. Photosynthetic types were marked with different colors, red: C_3_; green: C_3_-C_4_; purple: C_4_-like; blue: C_4_. (B) The transcript abundance of PPT1 and PPT2 in leaf and root. (C) The transcript abundance of PPT1 and PPT2 in BSC and whole leaf or MC in one C_3_ species (*Arabidopsis*) and four C_4_ species. All the data were from published RNA-SEQ data; the data source was detailed in the Method section. (Abbreviations: BSC: bundle sheath cell, WL: whole leaf, MC: mesophyll cell. Species abbreviation were listed in Table S1)

We then investigated whether the observed increase of transcript abundance for PPT1 is leaf-specific. To do this, we compared the transcript abundance of PPT1 and PPT2 in both root and leaf tissues in closely related C_3_ and C_4_ species using public RNA-SEQ data (Kulahoglu *et al.*, 2014; Moreno-Villena *et al.*, 2018) (Fig. 2 B). In leaf, PPT1 showed transcript abundance similar to PPT2 in C_3_ species but much higher transcript abundance than that of PPT2 in C_4_ species (Fig. 2B). However, the expression patterns of PPTs were drastically different in root. For the C_3_ dicot species, PPT1 usually showed higher transcript abundance than PPT2, the same pattern was found in most C_3_ species of monocot with *Lasiacis sorghoidea* (Lsor) an exception. In C_4_ monocot species, PPT1 showed no consistently higher expression abundance than PPT2 in root. Furthermore, the PPT expression levels in root were generally lower in C_4_ as compared to that in C_3_ species (Fig. 2 B).

After examining the evolutionary changes of tissue specificity of PPTs at the transcriptional level, we further investigated the cellular specificity of PPT expression. Specifically, we compared the transcript abundance of PPT1 and PPT2 in BSC and whole leaf in one C_3_ species and that of BSC and MC in four C_4_ species (Fig. 2 C). RNA-SEQ data from transcript residency on ribosome (Aubry *et al.*, 2014) show that PPT1 had higher expression level in BSC than in the whole leaf in *Arabidopsis*, whereas PPT2 displayed the opposite pattern, which is consistent with earlier histochemical localization of PPTs promoter (Knappe *et al.*, 2003). In all C_4_ species examined in this study, the transcript abundance of PPT1 was consistently higher in MC than in BSC (Fig. 2 C). In contrast, PPT2 showed an inconsistent pattern in the two cell types (Fig. 3C).

**Figure 3.**
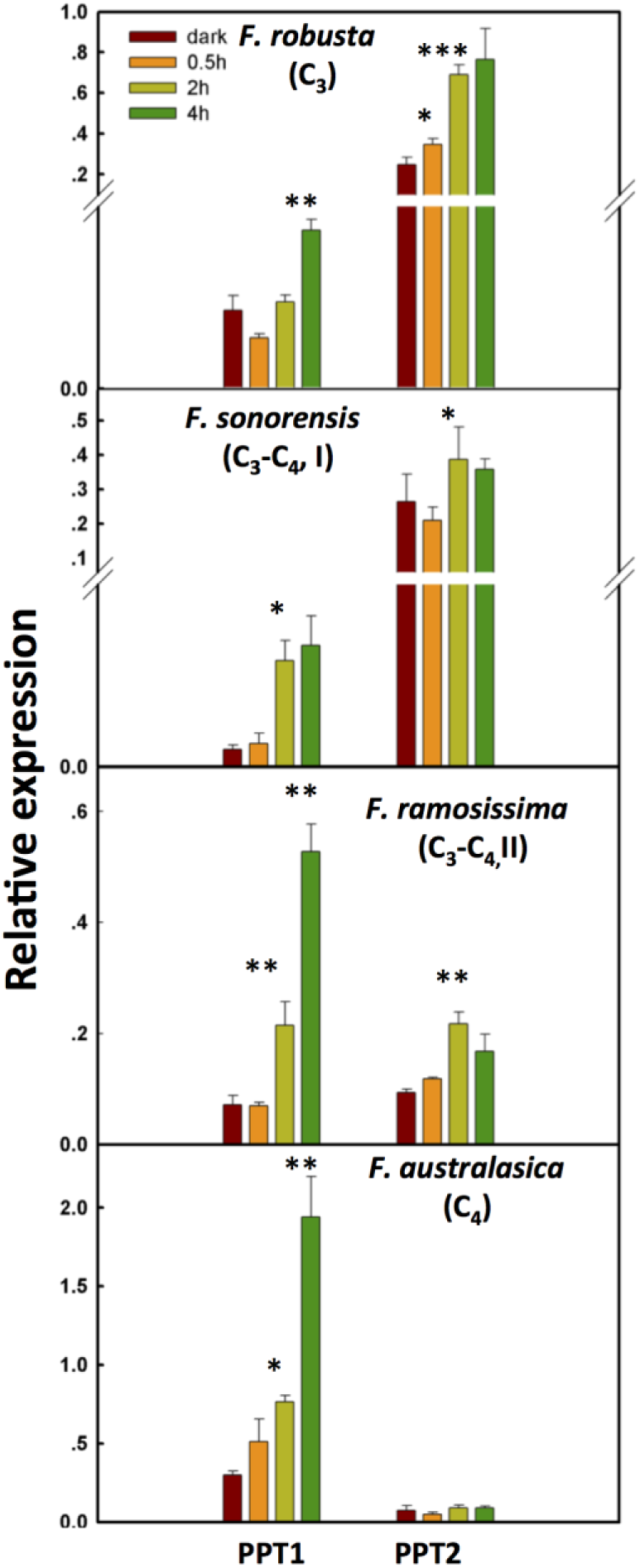
The change in light responsiveness of PPT1 and PPT2 along the evolution of C_4_ photosynthesis. qRT-PCR was used to quantify the transcript abundance of PPT1 and PPT2 in mature leaves after 0h, 0.5h, 2h and 4h after illumination. Significance levels represent the significance of the difference between the transcript abundance at a time point compared to that at the proceeding time point. (T-Test; *: 0.05~0.01, **: 0.01~0.001, ***: <0.001).

Taking together, along the evolution of C_4_ photosynthesis, PPT1 on one hand increased its expression abundance in leaf; on the other hand, PPT1 shifted its site of expression from dominantly in the BSC in C_3_ species to in MC in C_4_ species.

### The changes in light responsiveness of PPT during evolution from C_3_ to C_4_ species

How did PPT1 gained increased transcript abundance in leaf MC compared to its C_3_ ancestral PPT1? Given that the leaf MC typically receives more light than BSC (Xiao *et al.*, 2016), one possibility is that the C_4_ PPT1 might have acquired light-responsive *cis-*elements, which enables PPT1 to be more responsive to light induction. To test the possibility, we first investigated the light responsiveness of the two PPT paralogs along the C_4_ phylogeny. Specifically, we compared the transcript abundance of PPT1 and PPT2 in mature leaves after 0, 0.5h, 2h and 4h of illumination respectively. We quantified the transcript abundance using qRT-PCR in four *Flaveria* species, representing C_3_ photosynthesis, *F. robusta*, type I C_3_-C_4_ species *F. sonorensis*, type II C_3_-C_4_ species *F. ramosissima* and C_4_ species *F. australasica* (Fig. 3) (Sage *et al.*, 2012). Our results demonstrate a gradual increase in the speed of changes of PPT1 transcript abundance to light from C_3_ to C_3_-C_4_ intermediate to C_4_ species. Specifically, the transcript abundance of PPT1 did not show a significant up-regulation (P<0.05, T-Test) until 4h after illumination in the C_3_ *F. robusta*, whereas a significant up-regulation of PPT1 transcript abundance was observed at 2h in C_3_-C_4_ species. In the C_4_ species, the transcript abundance of PPT1 showed an up-regulation at 0.5 hour after illumination with a marginal significant level (P=0.075, T-Test).

We further examined whether the patterns of the enhancement in transcript abundance of PPT upon illumination changed along the evolution from C_3_ to C_4_ species. Type I C_3_-C_4_ species showed the maximal PPT1 transcript abundance at 2h after illumination, while the transcript abundance of PPT1 in type II C_3_-C_4_ and C_4_ species kept increasing even 4h after illumination (Fig. 3). Interestingly, the light response of PPT2 showed an opposite evolutionary pattern as compared to PPT1. Specifically, in the C_3_ *F. robusta*, PPT2 showed significantly higher transcript abundance than PPT1. An up-regulated expression level of PPT2 in *F. robusta* was observed at 0.5h after illumination, and further enhancement were observed till 2h. Nevertheless, in both C_3_-C_4_ species, the significantly enhanced expression of PPT2 was not detected until 2h after given light. In contrast, the transcript abundance of PPT2 showed no significant up-regulation in C_4_ species. Therefore, along the evolutionary trajectory, PPT1 gained not only higher transcript abundance in leaf, in particular in the MC, but also gained a more rapid and long-lasting response to light illumination, while PPT2 gradually lost its light responsiveness.

One parsimonious explanation to the increased leaf MC transcript abundance and also increased light responsiveness of PPT1 is that it may have recruited new light responsive *cis-*elements or trans-factors. To test this possibility, we predicted *cis*-elements on 3k base pairs (bp) upstream from the transcription start site of both PPT1 and PPT2 from four *Flaveria* species, namely, *F. robusta* (C_3_)*, F. sonorensis* (C_3_-C_4_)*, F. ramosissima* (C_3_-C_4_) and *F. trinervia* (C_4_), using the online tool Plantpan2.0 (http://plantpan2.itps.ncku.edu.tw/) with a score threshold of 0.9. We found that both PPT1 and PPT2 recruited new *cis*-elements at the transition between C_3_ to C_3_-C_4_ stage, e.g., CACTFTPPCA1: a MC specific *cis-*element detected in PEPC (Akyildiz *et al.*, 2007; Gowik *et al.*, 2004); CAATBOX1: a tissue specific enhancer (Muthamilarasan *et al.*, 2015); and GT1CONSENSUS (Shu *et al.*, 2015) and IBOXCORE (Martinez-Hernandez *et al.*, 2002), light responsive *cis-*elements (Table 1, Supplemental file 2). Therefore, we hypothesize that the recruitment of new *cis*-elements may underlie the observed changes in the expression abundance, expression location and its light responsiveness. It is worth noting here that in PPT2, new *cis-*elements were also recruited similarly even though PPT2 showed a different change with PPT1 in the responsiveness to light during C_4_ evolution (Table 1 and Fig. 3).

**Table 1.** *cis*-elements recruited into the promoter of PPT1 and PPPT2 during evolution

| *Cis*-elements | Frob (C<sub>3</sub>) |  | Fson (C<sub>3</sub>-C<sub>4</sub>) |  | Fram (C<sub>3</sub>-C<sub>4</sub>) |  | Ftri (C<sub>4</sub>) |  | Function | Reference |
| --- | --- | --- | --- | --- | --- | --- | --- | --- | --- | --- |
|  | PPT1 | PPT2 | PPT1 | PPT2 | PPT1 | PPT2 | PPT1 | PPT2 |  |  |
| DOFCOREZM | 0 | 51 | 45 | 47 | 41 | 53 | 32 | 56 | increase PPDK, decrease PEPC | Yanagisawa et al, 2000 |
| CACTFTPPCA1 | 0 | 0 | 45 | 48 | 38 | 33 | 43 | 39 | MC sepcific, leaf specific | Gowik et al., 2004 |
| CAATBOX1 | 0 | 0 | 37 | 43 | 41 | 43 | 37 | 36 | enhancer,tissue-specifict | Muthamilarasan et al., 2015 |
| IBOXCORE | 0 | 0 | 3 | 10 | 6 | 8 | 7 | 10 | light response | Kiseleva etal., 2017 |
| GT1CONSENSUS | 0 | 0 | 20 | 28 | 27 | 25 | 26 | 31 | light response | Michael and Davi, 1998 |
| ARR1AT | 0 | 0 | 37 | 31 | 27 | 31 | 26 | 32 | TF binding motif | Kim et al., 2010, |
| ROOTMOTIFTAPOX1 | 0 | 0 | 35 | 18 | 47 | 17 | 34 | 30 | tissue development,root specific | Elmayan and Tepfer, 1995 |
| GTGANTG10 | 0 | 0 | 19 | 26 | 21 | 25 | 25 | 24 | tissue development | Shu et al., 2015 |
| POLLEN1LELAT52 | 0 | 0 | 18 | 22 | 18 | 21 | 11 | 15 | tissue development | Borges et al., 2011 |
| TAAAGSTKST1 | 0 | 0 | 14 | 15 | 11 | 13 | 9 | 13 | Dof specific | venkatesh et al., 2017 |
| SEF4MOTIFGM7S | 0 | 0 | 21 | 5 | 21 | 4 | 18 | 11 | seed and embryo specific expression | Kaur et al., 2015 |

Note: The values show the copy numbers of *cis*-elements. (Abbreviations: Frob: *F. robusta*; Fson: *F. sonorensis*; Fram: *F. ramosissima*; Ftri: *F. trinervia*).

### Amino acid changes in PPT during the evolution from C_3_ to C_4_ species

Besides changes in the expression patterns and transcript abundance, another factor that contributes to gene neofunctionalization is accumulation of mutations supporting the new function (Christin *et al.*, 2013). To test whether this is the case in PPT, we compared amino acid sequence of PPT1 and PPT2 for 16 species in the genus *Flaveria*, covering C_3_, C_3_-C_4_, C_4_-like and C_4_ species. The results show that, PPT1 has more consistent amino acid modifications than PPT2 when the sequences between C_4_ and C_3_ species were compared. The consistent amino acid modifications are defined as those amino acid sequence which are consistent in C_4_ species but different with that in C_3_ species. Specifically, amino acid sequence of PPT1 has 19 consistent amino acid modifications between C_3_ and C_4_ species, while in contrast, PPT2 exhibits 8 consistent amino acid modifications (Fig. 4). Interestingly, in this analysis, PPT1 shows no signal of positive selection in C_4_ species when compared to C_3_ species in the genus of *Flaveria*. PPT2 in contrast shows a signal of positive selection in C_4_ species in the same genus; however, none of the two predicted positive selected sites of PPT2 is neither C_4_ specific nor C_4_ consistent modifications (Fig. S1). Interestingly, PPT1 accumulated a large segment of insertion which emerged at the common ancestor of C_4_-like and C_4_ species in Clade A. The insertion segments had either four (*F. pameri*, *F. bidentis*, *F. trinervia* and *F. australasica*) or five (*F. vaginata* and *F. konchiana*) repeats with each repeat comprising 13 amino acids, *i.e.*, 13-aa element (Fig. S2). And the number of 13-aa elements is conserved at different development stage in those species (Fig. S2).

**Figure 4.**
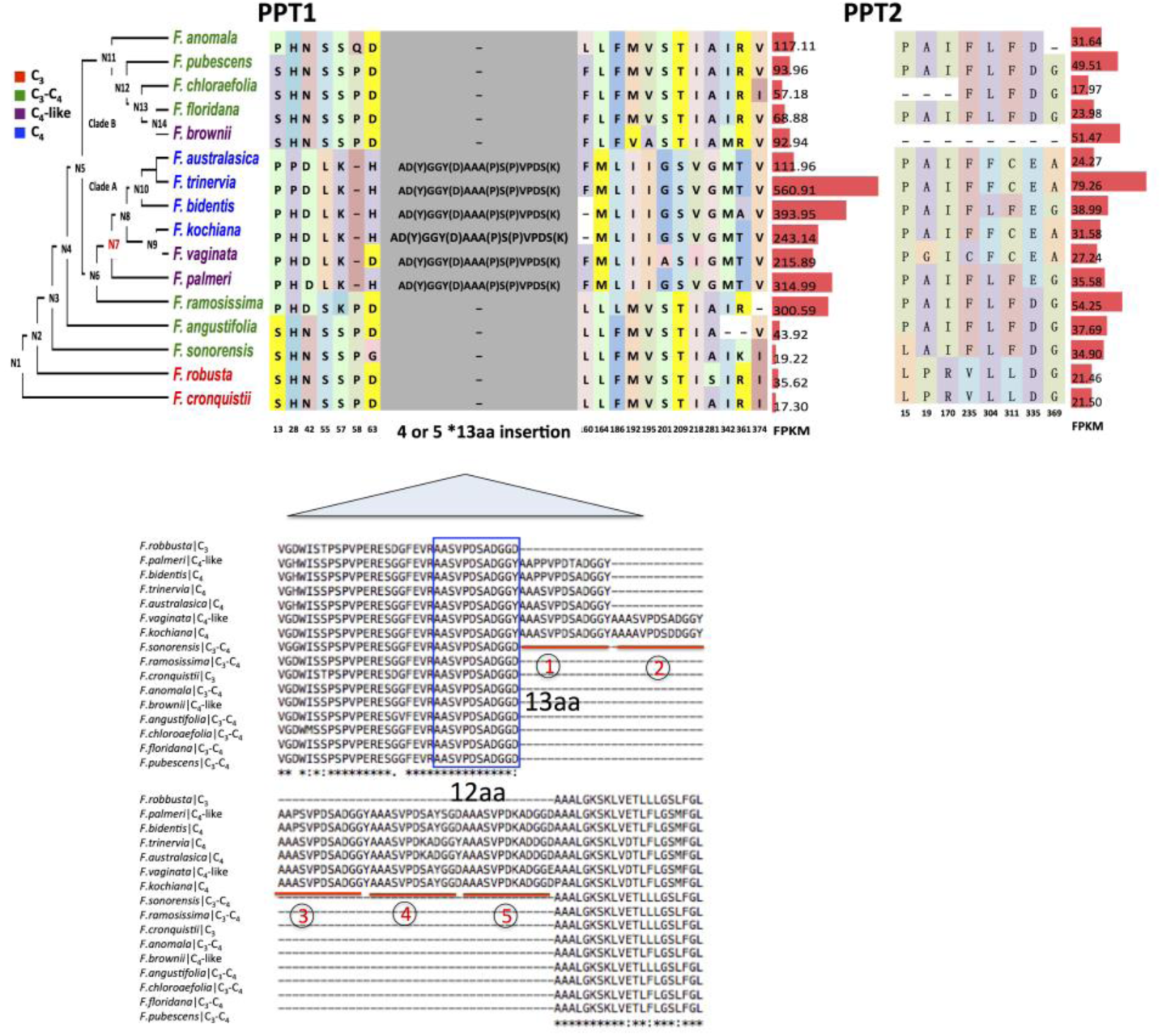
The evolution of PPT1 and PPT2 protein sequences in the genus *Flaveria*. Amino acid changes of PPT1 and PPT2 comparing C_4_ to C_3_ species are showed on the trace of the phylogeny. Transcript abundance calculated as FPKM was displayed on the right with red bars. Numbers below amino acids are the aligned locations. Symbols “-” presenting alignment gap. PPT1 (406 amino acid in *F. cronquistii*) showed more frequently amino acid changes than PPT2 (417 amino acid in *F. cronquistii*). An insertion composed of four or five 13-aa elements occurred at the ancestral node “N7” (marked in red) on the phylogenetic tree. The sequence of the 13-aa segment is variant AAA(P)SVPDS(K)AD(Y)GGY(D) at four sites.

### PPT1 from C_4_ species is a bidirectional transport of PEP

The metabolic processes of C_3_ and C_4_ photosynthesis require change of the direction of PEP transport between cytosol and chloroplast stroma. So far, there is no explicit study on the direction of PEP transport in C_4_ species. We hypothesized that in C_4_ photosynthesis, the C_4_ version PEP transporter, i.e. PPT1, might specifically transfer PEP from chloroplast to cytosol. It is possible that the extra insertion segment emerged during C_4_ evolution might be related to this direction of transport. We tested this hypothesis with a genetic approach by expressing *Flaveria* PPT1 in a C_3_ *Arabidopsis* PPT1 loss-of-function mutant *cue1-5* (Li *et al.*, 1995). We predicted that a PPT1 importing PEP from cytosol into chloroplast should rescue the phenotype *cue1-5* mutant while a PPT1 exporting PEP from chloroplast to cytosol should not rescue this phenotype. We hence generated C-terminal GFP fused PPT1 from four different *Flaveria* species, including one C_3_ species *F. cronquistii*, two intermediate species *F. ramosissima* (C_3_-C_4_) and *F. plameri* (C_4_-like), and one C_4_ species *F. bidentis* (C_4_), and expressed these PPT1-GFP driven by a *35S* promoter in *cue1-5* through a transgenic method (Fig. 5A). The result shows that PPT1 from all four species complemented the reticulate leaf phenotype and small rosette size of *cue1-5* (Fig. 5B and 5C). This data not only indicate PPT1 is functionally conserved in these different *Flaveria* species and also *Arabidopsis*, but also suggest that PPT1 from C_4_ *Flaveria* C_4_ can import PEP from cytosol into chloroplast as well. In another word, our data suggest that C_4_ PPT1 is a bi-directional PEP transporter.

**Figure 5.**
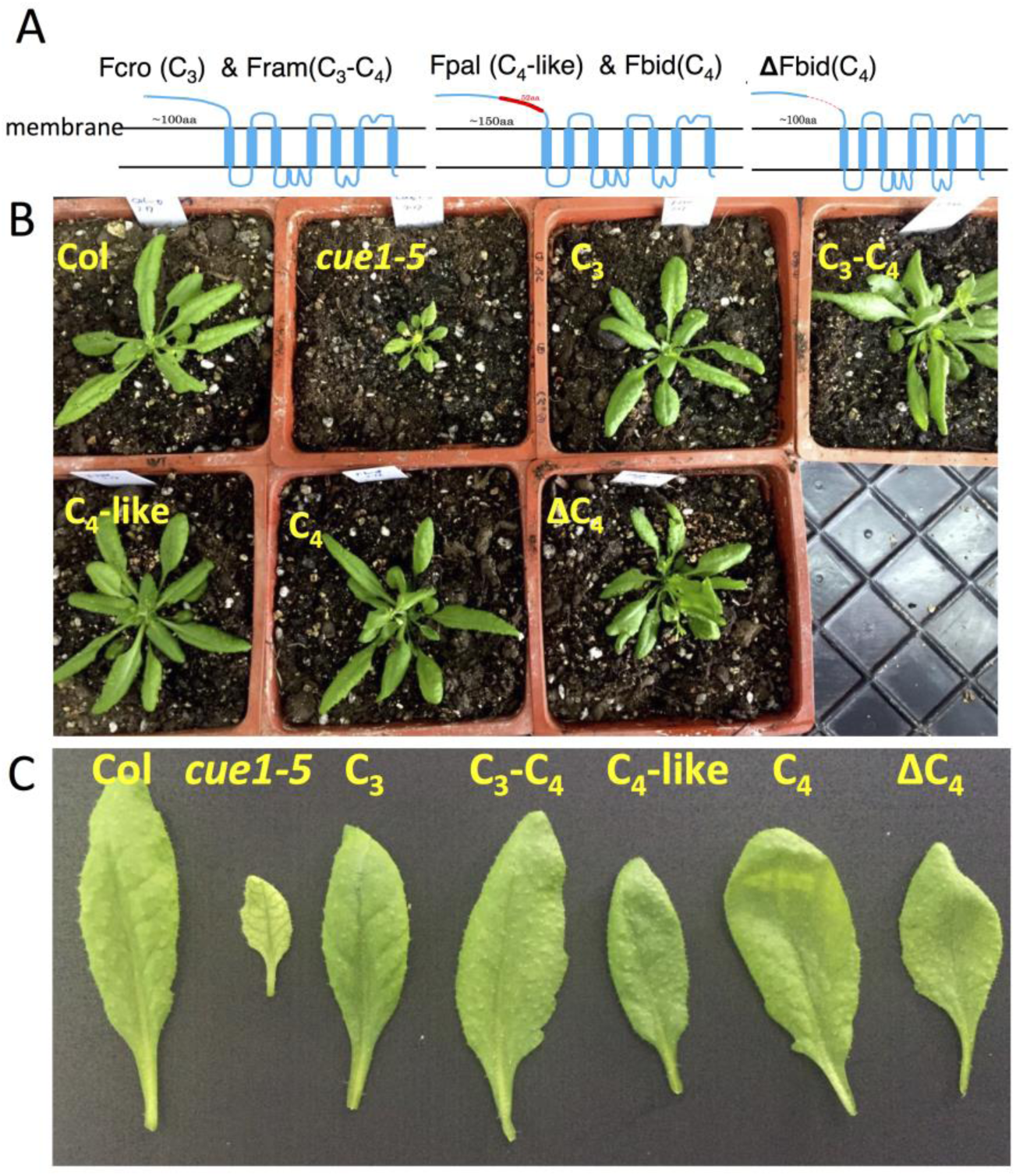
*Flaveria* PPT1 complements the phenotype of *Arabidopsis* PPT1 lose-of-function mutant *cue1-5*. (A) Shows the position of 52-aa insertion in protein sequence of *F. bidentis* (C_4_) PPT1. The insertion was predicted to be located at the non-membrane-portion. (B) and (C) *Arabidopsis cue1-5* shows reticulate leaf phenotype and decreased rosette size. PPT1 from different types of photosynthetic *Flaveria* species rescued the phenotype of *Arabidopsis cue1-5*. Besides, PPT1 from *F. bidentis* without the insertion (ΔC_4_) also recovers the phenotype of the *Arabidopsis cue1-5*. (Abbreviations: Fcro, *F. cronquistii*, Fram: *F. ramosissima*, Fpal: *F. palmeri*, Fbid: *F. bidentis*.)

Even if this extra insertion segment has no influence on metabolite transport direction, may it have other physiological functions? We explored this question by removing the 4×13-aa insertion in *F. bidentis* (C_4_) (ΔFbidPPT1 for short) and expressing it in *cue1-5 Arabidopsis* (Fig. 5A). The transgenic plant *ΔFbidPPT1*/*cue1* showed the same phenotype as *FbidPPT1*/*cue1* (Figs. 5B and 5C), suggesting that the insertion does not affect the PEP transport function of PPT1. It is likely therefore this extra insertion does not influence the structure of this protein in thylakoid membrane. Indeed, protein structure prediction using I-TASSER shows that the insertion site lies in the outer membrane portion of FbidPPT1 (Fig. 5A), which possibly does not influence the structure of PPT1 in thylakoid membrane. We further checked whether the extra insertion affects the subcellular location of PPT1 by using transient expression of PPT1-GFP in *N. benthamiana* leaves. Both the FbidPPT1 and ΔFbidPPT1 were shown to localize in chloroplast (Figs. S3A and S3C), suggesting the insertion has no impact on the subcellular localization of PPT1 as well. Further experiments show that PPT1 and PPT2 from all four types of *Flaveria* species were localized to chloroplast (Fig. S3).

## Discussion

PPT1 experienced various of changes during its recruitment into C_4_ photosynthesis. First, PPT1 transcript abundance was lower than PPT2 in C_3_ species, while in C_4_ species its transcript abundance became much higher than PPT2. Secondly, its expression location shifted from predominantly in roots in C_3_ plants to predominantly in leaves in C_4_ plants. Thirdly, it expressed preferably in BSC in C_3_ plants while preferably in MC C_4_ plants. Fourthly, it gained speedy and long-lasting responsiveness to light in C_4_ compared to C_3_ plants. Fifthly, it accumulated many *cis*-elements and new amino acid modifications during the evolution of C_4_ photosynthesis. In this session, we will briefly discuss these in the context of C_4_ evolution.

### Using a parsimonious solution: a recurring theme to solve incident challenges during the C_4_ PPT1 evolution

A recurring theme during C_4_ evolution is the extensive re-utilization of pre-existing components, including metabolic genes (Aubry *et al.*, 2011), *cis-*regulatory elements (Brown *et al.*, 2011; Kajala *et al.*, 2012), signaling pathways (Burgess *et al.*, 2016a), and anatomical features (Griffiths *et al.*, 2013). During the transition from C_3_ to C_4_ photosynthesis, usually the C_3_ ancestral copy of a gene acquired new cellular specificity for its expression, altered its expression abundance and sometimes acquired new sequence variations conferring adaptive advantage (Hibberd and Covshoff, 2010; Paulus *et al.*, 2013). Here we show that during the evolution of C_4_ photosynthesis, C_4_ PPT1 altered its expression location from predominantly in root to leaf, and from BSC to MC, furthermore, it greatly increased its transcript abundance and responsiveness to light (Figs. 2 and 3). Meanwhile, PPT1 gained many new *cis-*elements at the promoter region (3k bp upstream of the start codon). The function of these *cis-*elements ranges from elements responsive to light (GT1CONSENSUS (Shu *et al.*, 2015)) to elements controlling tissue specificity (CAATBOX1 (Muthamilarasan *et al.*, 2015)) and elements controlling cell specificity (CACTFTPPCA1 (Akyildiz *et al.*, 2007; Gowik *et al.*, 2004), DOFCOREZM (Yanagisawa, 2000)). Interestingly, all these *cis*-elements that were predicted to be newly recruited into the promoter of PPT1 during C_4_ evolution were also predicted to exist in promoters of other genes in the C_4_ shuttle (including NADP-ME, PEPC, PEPC-k, PPDK and PPDK-RP), H and P subunit of Glycine decarboxylase (Supplemental file 2). It is worth emphasizing here molecular experiments are needed to study the functional significance of these *cis-*elements.

Interestingly, these different *cis*-elements were predicted to be in many C_4_ shuttle genes in the C_3_-C_4_ type I intermediate species *F. sonorensis* (Table 1), implying potential adaptive advantage of gaining these elements simultaneously compared to an alternative case where elements were recruited gradually. Notably, the *cis-*element ODFCOREZM also resided in the promoters of PEPC (Gowik *et al.*, 2004; Yanagisawa, 2000) and PPDK (Table 1, Supplemental file 2). This is probably not surprising given that the PPDK catalyzes generation of PEP, while PEPC uses PEP as its substrate. Possessing a common *cis-*element in PPT1, PEPC and PPDK enables coordinated expression of these different proteins. It’s worth emphasizing here that these potential *cis-*elements were identified through sequence comparison, their roles in controlling the expression patterns of the PPT1 need further experimental verifications. Interestingly, our analysis of the promoter sequence suggest that PPT2 also acquired the same *cis-*elements as PPT1, however, PPT2 gained neither increased expression abundance, nor preferred leaf or MC expression in C_4_ species (Table 1, Figs. 2, 3).

The direction of PEP transport between chloroplast and cytosol in MC should be reversed between C_3_ and C_4_ photosynthesis theoretically. In C_3_ mesophyll cells, PEP needs to be imported into chloroplast to provide intermediate for shikimate pathway, however, PEP needs to be exported from chloroplast stroma to cytosol in C_4_ MC (Weber and Linka, 2011). Given this, we expect that the direction of PPT1 in the C_4_ species might be changed from that in C_3_ species. However, our results show that PPT1 from C_4_ species fully complemented the leaf phenotype of a C_3_ *Arabidopsis* PPT1 loss-of-function mutant *cue1-5* (Fig. 5), suggesting that PPT1 is a bi-directional PEP transporter. Having a bi-directional PPT1 transporter may enable plants have higher adaptability since it would allow the metabolic system to adapt to different metabolic situations. Again, this may be related to the tendency of plants to utilize the most parsimonious solution to solve incident challenges. On one side, during the C_4_ photosynthesis, if the concentration of PEP in chloroplast is not higher than that in cytosol, there will be no metabolic demand for transport of PEP from chloroplast to cytosol; on the other side, once a concentration gradient of PEP cross thylakoid membrane is established, there is no demand for a uniporter of PEP. In another word, the bi-directionality of PPT1 is again consistent with the principle of taking an easy route to acquire a new function for an existing protein.

### Less abundant PPT in ancestral C_3_: a metabolic pre-condition for C_4_ evolution?

According to the current theory of C_4_ evolution, the shift of expression location of glycine decarboxylase (GDC) from MC to BSC may bridge the evolution from C_3_ to C_4_ photosynthesis (Mallmann *et al.*, 2014). The decarboxylation of glycine by GDC creates nitrogen imbalance between BSC and MC, which necessitates the formation of nitrogen balancing pathways (Mallmann *et al.*, 2014). Different pathways were proposed, among which one has PPT1 involved and the direction of the PEP transport in this pathway is from chloroplast to cytosol; this is the pathway which can later be optimized to evolve C_4_ photosynthetic pathway (Mallmann *et al.*, 2014). Considering that this hypothetical ancestral pathway needs PEP export from chloroplast to cytosol, which is in contrast to the direction of PEP transport in a typical C_3_ plant, we hypothesize that C_3_ plants which have relatively less capacity or demand for import of PEP from cytosol to chloroplast might be easier to adapt its metabolism to a new metabolic state.

Consistent with this hypothesis, the C_3_ species in genera shown in Fig. 2A, which represent genera with higher propensity of evolving C_4_, showed lower total PPT transcript abundance as compared to that in *Arabidopsis*, which is in the Brassica genus with no recorded C_4_ species (Fig. 2). Furthermore, our earlier study shows that the expression patterns of PPT1 and PPT2 differ between C_3_ species from genera that with abundant C_4_ species (C_4_-rich C_3_ species) and C_3_ species from genera with bare C_4_ species (C_4_-poor C_3_ species) (Tao *et al.*, 2016). Specifically, PPT2 showed higher transcript abundance than PPT1 in C_4_-poor C_3_ species while the scenario is reverse in C_4_-rich C_3_ species. Moreover, C_4_-rich C_3_ species have lower total transcript abundance of PPT than C_4_-poor C_3_ species (*P* <0.05, T-Test) (Fig. S4). This observation suggests that the evolution of C_4_ photosynthetic metabolism might, to certain degree, be connected to the changes in the PEP metabolism, i.e. the production, transport and consumption of PEP in C_3_ species. Having overall lower PPT transport but relatively higher PPT1 expression might be a metabolic pre-condition for evolution of C_4_ photosynthetic metabolism. More data of PPT transcript abundance in additional C_4_-rich C_3_ species and C_4_-poor C_3_ species are needs to further test this hypothesis.

Is there a physiological basis for a linkage between PPT abundance and the evolvability of C_4_ photosynthesis? PEP is a starting substrate used in the shikimate pathway. The shikimate pathway produces chorismite, which is the basis for a large array of secondary metabolites, including alkaloids, flavonoids, and lignin; many of these are important players in plant defense mechanisms and stress responses (Herrmann, 1995; Iriti and Faoro, 2009). Having higher PPT activity may enable plants to produce higher amount of secondary metabolites, hence conferring plants with enhanced tolerance to stresses. From this perspective, C_3_ plants with higher PPT activity may have increased stress tolerance and experienced less evolutionary pressure to evolve C_4_ under severe stresses. In another word, evolving a C_4_ mechanism may be only one of potential solutions plants use to cope with stress conditions promoting evolution of C_4_ photosynthesis, such as drought, salinity stress, and high temperature stress etc. (Osborne and Freckleton, 2009; Sage, 2003). This notion is consistent with the common observation that even in extremely arid conditions, such as desert, C_4_ and C_3_ plants co-exist (Li *et al.*, 2018; Su *et al.*, 2012) demonstrating the natural diversity of options to cope with the arid stress.

### Changes in the amino acid sequences in PPT1 during the evolution from C_3_ to C_4_ photosynthesis

Besides having much higher transcript abundance, expression patterns and responsiveness to light, PPT1 also gained many amino acid modifications during its transition to C_4_ version (Fig. 4). PPT1 has more amino acid modifications during the evolution than PPT2, but none of these modifications showed positive selection signal in C_4_ species. This is consistent with the situation for other genes, such as PEPC, PPDK, and NAD-ME reported in our earlier study (Lyu *et al.*, 2018). Interestingly, PPT2 though showed positive selection in *Flaveria* C_4_ species compared to C_3_ *Flaveria* species, but it does not show positive selection in C_4_ grass species compared to C_3_ grass species (Huang *et al.*, 2017), suggesting that PPT2 in different plant lineages might have experienced different evolutionary trajectories.

In addition to the individual single amino acid changes, our results also show a four or five 13-aa elements insertion in PPT1 of C_4_-like and C_4_ *Flaveria* species in the clade A from the phylogenetic tree (Fig. 4). This extra insertion neither influences the location of expression of PPT1 (Fig. 4), nor the capacity of PPT1 to complement the phenotype of *Arabidopsis cue1-5* mutant (Fig. 5). Furthermore, we did not find this insertion in PPTs from other species (Fig. S5). The potential physiological significance of this insertion still needs to be explored, if there is any. Here the lack of physiological role of the identified insertion of these 13-aa units into PPT1 during C_4_ evolution highlights the need of molecular experiments to confirm any conclusion derived from sequence analysis.

## Author contribution

XGZ, GC and MJL designed the project and wrote the paper, MJL did bioinformatics analysis and qRT-PCR, YW and JJ conducted the transgenic experiments.

## Acknowledgement

The authors appreciate Prof. Rowan Sage and Prof. Peter Westhoff for sharing us *Flaveria* materials. We thank Prof. Daiyin Chao and Prof. Haiyang Hu for great discussion and suggestion; MJL also thanks Faming Chen, Fengfeng Miao and Yongyao Zhao for help in experiment parts. This work was sponsored by Shanghai Sailing Program [17YF421900], National Science Foundation of China [31701139] and [31500988].

## Supplementary data

Supplemental file 1: supplemental figures and tables

Figure S1. Two amino acid showing signal of positive selection of PPT2 in C_4_ species

Figure S2. The insertion of 13aa-element in C_4_-like and C_4_ PPT1

Figure S3. Subcellular localization of PPT1 and PPT2 in *Flaveria* species

Figure S4. The transcript abundance of PPT in C_3_ species from C_4_ poor and C_4_ rich genera

Figure S5. The 13-aa-element insertion is not universal.

Table S1. Abbreviations of species used in this study

Table S2. Primers used in this study

Supplemental file 2: predicted *cis*-elements of PPT and other genes from the promoter sequence.

## References

Akyildiz M, Gowik U, Engelmann S, Koczor M, Streubel M, Westhoff P. 2007. Evolution and function of a cis-regulatory module for mesophyll-specific gene expression in the C_4_ dicot Flaveria trinervia. Plant Cell 19: 3391–3402.

Aubry S, Brown NJ, Hibberd JM. 2011. The role of proteins in C_3_ plants prior to their recruitment into the C_4_ pathway. Journal of Experimental Botany 62: 3049–3059.

Aubry S, Smith-Unna RD, Boursnell CM, Kopriva S, Hibberd JM. 2014. Transcript residency on ribosomes reveals a key role for the Arabidopsis thaliana bundle sheath in sulfur and glucosinolate metabolism. Plant Journal 78: 659–673.

Bailey KJ, Leegood RC. 2016. Nitrogen recycling from the xylem in rice leaves: dependence upon metabolism and associated changes in xylem hydraulics. Journal of Experimental Botany 67: 2901–2911.

Brown NJ, Newell CA, Stanley S, Chen JE, Perrin AJ, Kajala K, Hibberd JM. 2011. Independent and parallel recruitment of preexisting mechanisms underlying C_4_ photosynthesis. Science 331: 1436–1439.

Burgess SJ, Granero-Moya I, Grange-Guermente MJ, Boursnell C, Terry MJ, Hibberd JM. 2016. Ancestral light and chloroplast regulation form the foundations for C_4_ gene expression. Nature Plants 2: 16161.

Camacho C, Coulouris G, Avagyan V, Ma N, Papadopoulos J, Bealer K, Madden TL. 2009. BLAST+: architecture and applications. BMC Bioinformatics 10: 421.

Chang YM, Liu WY, Shih AC, et al. 2012. Characterizing regulatory and functional differentiation between maize mesophyll and bundle sheath cells by transcriptomic analysis. Plant Physiology 160: 165–177.

Christin PA, Besnard G. 2009. Two independent C_4_ origins in Aristidoideae (Poaceae) revealed by the recruitment of distinct phosphoenolpyruvate carboxylase genes. American Journal of Botany 96: 2234–2239.

Christin PA, Boxall SF, Gregory R, Edwards EJ, Hartwell J, Osborne CP. 2013. Parallel recruitment of multiple genes into C_4_ photosynthesis. Genome Biology and Evolution 5: 2174–2187.

Christin PA, Petitpierre B, Salamin N, Buchi L, Besnard G. 2009. Evolution of C_4_ phosphoenolpyruvate carboxykinase in Grasses, from genotype to phenotype. Molecular Biology and Evolution 26: 357–365.

Denton AK, Mass J, Kulahoglu C, Lercher MJ, Brautigam A, Weber AP. 2017. Freeze-quenched maize mesophyll and bundle sheath separation uncovers bias in previous tissue-specific RNA-Seq data. Journal of Experimental Botany 68: 147–160.

Edgar RC. 2004. MUSCLE: multiple sequence alignment with high accuracy and high throughput. Nucleic Acids Research 32: 1792–1797.

Emms DM, Covshoff S, Hibberd JM, Kelly S. 2016. Independent and parallel evolution of new genes by gene duplication in two origins of C_4_ photosynthesis provides new insight into the mechanism of phloem loading in C_4_ species. Molecular Biology and Evolution 33: 1796–1806.

Fischer K, Kammerer B, Gutensohn M, Arbinger B, Weber A, Hausler RE, Flugge UI. 1997. A new class of plastidic phosphate translocators: A putative link between primary and secondary metabolism by the phosphoenolpyruvate/phosphate antiporter. Plant Cell 9: 453–462.

Gowik U, Burscheidt J, Akyildiz M, Schlue U, Koczor M, Streubel M, Westhoff P. 2004. cis-Regulatory elements for mesophyll-specific gene expression in the C_4_ plant Flaveria trinervia, the promoter of the C_4_ phosphoenolpyruvate carboxylase gene. Plant Cell 16: 1077–1090.

Griffiths H, Weller G, Toy LFM, Dennis RJ. 2013. You’re so vein: bundle sheath physiology, phylogeny and evolution in C_3_ and C_4_ plants. Plant Cell and Environment 36: 249–261.

Hatch MD. 1987. C_4_ photosynthesis – a unique blend of modified biochemistry, anatomy and ultrastructure. Biochimica et Biophysica Acta 895: 81–106.

Hatch MD, Osmond CB. 1976. Compartmentation and transport in C_4_ photosynthesis. Encyclopedia of Plant Physiology 3: 144–184.

Herrmann KM. 1995. The shikimate pathway – early seps in the biosynthesis of aromatic-compounds. Plant Cell 7: 907–919.

Herrmann KM, Weaver LM. 1999. The shikimate pathway. Annual Reveiw of Plant Biology 50: 473–503.

Hibberd JM, Covshoff S. 2010. The regulation of gene expression required for C_4_ photosynthesis. Annual Review of Plant Biology 61: 181–207.

Huang P, Studer AJ, Schnable JC, Kellogg EA, Brutnell TP. 2017. Cross species selection scans identify components of C_4_ photosynthesis in the grasses. Journal of Experimental Botany 68: 127–135.

Iriti M, Faoro F. 2009. Chemical diversity and defence metabolism: how plants cope with pathogens and ozone pollution. International Journal of Molecular Sciences 10: 3371–3399.

Jiao Y, Li J, Tang H, Paterson AH. 2014. Integrated syntenic and phylogenomic analyses reveal an ancient genome duplication in monocots. Plant Cell 26: 2792–2802.

John CR, Smith-Unna RD, Woodfield H, Covshoff S, Hibberd JM. 2014. Evolutionary convergence of cell-specific gene expression in independent lineages of C_4_ grasses. Plant Physiology 165: 62–75.

Kajala K, Brown NJ, Williams BP, Borrill P, Taylor LE, Hibberd JM. 2012. Multiple Arabidopsis genes primed for recruitment into C_4_ photosynthesis. Plant Journal 69: 47–56.

Knappe S, Lottgert T, Schneider A, Voll L, Flugge UI, Fischer K. 2003. Characterization of two functional phosphoenolpyruvate/phosphate translocator (PPT) genes in Arabidopsis--AtPPT1 may be involved in the provision of signals for correct mesophyll development. Plant Journal 36: 411–420.

Krebs HA. 1982. The evolution of the citric-acid cycle and other cyclic metabolic pathways. Chemiker-Zeitung 106: 86–87.

Kulahoglu C, Denton AK, Sommer M, et al. 2014. Comparative transcriptome atlases reveal altered gene expression modules between two Cleomaceae C_3_ and C_4_ plant species. Plant Cell 26: 3243–3260.

Li HM, Culligan K, Dixon RA, Chory J. 1995. Cue1 – a mesophyll cell-specific positive regulator of light-controlled gene-expression in Arabidopsis. Plant Cell 7: 1599–1610.

Li P, Ponnala L, Gandotra N, et al. 2010. The developmental dynamics of the maize leaf transcriptome. Nature Genetics 42: 1060–1067.

Li SJ, Su PX, Zhang HN, Zhou ZJ, Xie TT, Shi R, Gou W. 2018. Distribution patterns of desert plant diversity and relationship to soil properties in the Heihe River Basin, China. Ecosphere 9.

Lyu M-JA, Gowik U, Westhoff P, et al. 2018. The Coordination and Jumps along C_4_ Photosynthesis Evolution in the Genus Flaveria. bioRxiv: 460287.

Lyu MJ, Gowik U, Kelly S, et al. 2015. RNA-Seq based phylogeny recapitulates previous phylogeny of the genus Flaveria (Asteraceae) with some modifications. BMC Evolutionary Biology 15: 116.

Mallmann J, Heckmann D, Brautigam A, Lercher MJ, Weber AP, Westhoff P, Gowik U. 2014. The role of photorespiration during the evolution of C_4_ photosynthesis in the genus Flaveria. eLife 3: e02478.

Martinez-Hernandez A, Lopez-Ochoa L, Arguello-Astorga G, Herrera-Estrella L. 2002. Functional properties and regulatory complexity of a minimal RBCS light-responsive unit activated by phytochrome, cryptochrome, and plastid signals. Plant Physiology 128: 1223–1233.

Matasci N, Hung LH, Yan Z, et al. 2014. Data access for the 1,000 Plants (1KP) project. Gigascience 3: 17.

Moreno-Villena JJ, Dunning LT, Osborne CP, Christin PA. 2018. Highly expressed genes are preferentially co-opted for C_4_ photosynthesis. Molecular Biology and Evolution 35: 94–106.

Muthamilarasan M, Bonthala VS, Khandelwal R, Jaishankar J, Shweta S, Nawaz K, Prasad M. 2015. Global analysis of WRKY transcription factor superfamily in Setaria identifies potential candidates involved in abiotic stress signaling. Frontiers in Plant Science 6.

Osborne CP, Freckleton RP. 2009. Ecological selection pressures for C_4_ photosynthesis in the grasses. Proceedings of the Royal Society B: Biological Sciences 276: 1753–1760.

Paulus JK, Schlieper D, Groth G. 2013. Greater efficiency of photosynthetic carbon fixation due to single amino-acid substitution. Nature Communications 4: 1518.

Rao X, Lu N, Li G, Nakashima J, Tang Y, Dixon RA. 2016. Comparative cell-specific transcriptomics reveals differentiation of C_4_ photosynthesis pathways in switchgrass and other C_4_ lineages. Journal of Experimental Botany 67: 1649–1662.

Reyna-Llorens I, Hibberd JM. 2017. Recruitment of pre-existing networks during the evolution of C_4_ photosynthesis. Philosophical Transactions of the Royal Society B-Biological Sciences 372.

Sage RF. 2003. The evolution of C_4_ photosynthesis. New Phytologist 161.

Sage RF, Sage TL, Kocacinar F. 2012. Photorespiration and evolution of C_4_ photosynthesis. Annual Review of Plant Biology 63: 19–47.

Sage RF, Zhu XG. 2011. Exploiting the engine of C_4_ photosynthesis. Journal of Experimental Botany 62: 2989–3000.

Shu YJ, Song LL, Zhang J, Liu Y, Guo CH. 2015. Genome-wide identification and characterization of the Dof gene family in Medicago truncatula. Genetics and Molecular Research 14: 10645–10657.

Stamatakis A. 2006. RAxML-VI-HPC: Maximum likelihood-based phylogenetic analyses with thousands of taxa and mixed models. Bioinformatics 22: 2688–2690.

Stata M, Sage TL, Hoffmann N, Covshoff S, Ka-Shu Wong G, Sage RF. 2016. Mesophyll Chloroplast Investment in C_3_, C_4_ and C_2_ Species of the Genus *Flaveria*. Plant Cell Physiology 57: 904–918.

Su PX, Yan QD, Xie TT, Zhou ZJ, Gao S. 2012. Associated growth of C_3_ and C_4_ desert plants helps the C_3_ species at the cost of the C_4_ species. Acta Physiologiae Plantarum 34: 2057–2068.

Tao Y, Lyu MJ, Zhu XG. 2016. Transcriptome comparisons shed light on the pre-condition and potential barrier for C_4_ photosynthesis evolution in eudicots. Plant Molecular Biology 91: 193–209.

Tausta SL, Li P, Si Y, Gandotra N, Liu P, Sun Q, Brutnell TP, Nelson T. 2014. Developmental dynamics of Kranz cell transcriptional specificity in maize leaf reveals early onset of C_4_-related processes. Jouranl of Experimental Botany 65: 3543–3555.

Taylor L, Nunes-Nesi A, Parsley K, Leiss A, Leach G, Coates S, Wingler A, Fernie AR, Hibberd JM. 2010. Cytosolic pyruvate,orthophosphate dikinase functions in nitrogen remobilization during leaf senescence and limits individual seed growth and nitrogen content. Plant Journal 62: 641–652.

von Caemmerer S, Furbank RT. 2003. The C_4_ pathway: an efficient CO_2_ pump. Photosynthesis Research 77: 191–207.

Weber AP, von Caemmerer S. 2010. Plastid transport and metabolism of C_3_ and C_4_ plants--comparative analysis and possible biotechnological exploitation. Current Opinion in Plant Biology 13: 257–265.

Weber APM, Linka N. 2011. Connecting the plastid: transporters of the plastid envelope and their role in linking plastidial with cytosolic metabolism. Annual Review of Plant Biology 62: 53–77.

West-Eberhard MJ, Smith JAC, Winter K. 2011. Photosynthesis, reorganized. Science 332: 311–312.

Xiao Y, Tholen D, Zhu XG. 2016. The influence of leaf anatomy on the internal light environment and photosynthetic electron transport rate: exploration with a new leaf ray tracing model. Journal of Experimental Botany 67: 6021–6035.

Yanagisawa S. 2000. Dof1 and Dof2 transcription factors are associated with expression of multiple genes involved in carbon metabolism in maize. Plant Journal 21: 281–288.

Yang J, Zhang Y. 2015. I-TASSER server: new development for protein structure and function predictions. Nucleic Acids Research 43: W174–181.

Yang ZH. 2007. PAML 4: Phylogenetic analysis by maximum likelihood. Molecular Biology and Evolution 24: 1586–1591.

